# Coevolution of male and female reproductive traits drive cascading reinforcement in *Drosophila yakuba*

**DOI:** 10.1101/022244

**Authors:** Aaron A. Comeault, Aarti Venkat, Daniel R. Matute

**Author notes:** Correspondence: Biology Department, University of North Carolina, 250 Bell Tower Road, Chapel Hill, 27599.

## Abstract

When the ranges of two hybridizing species overlap, individuals may ‘waste’ gametes on inviable or infertile hybrids. In these cases, selection against maladaptive hybridization can lead to the evolution of enhanced reproductive isolation in a process called reinforcement. On the slopes of the African island of São Tomé, *Drosophila yakuba* and its endemic sister species *D. santomea* have a well-defined hybrid zone. *Drosophila yakuba* females from within this zone show increased postmating-prezygotic isolation towards *D. santomea* males when compared with *D. yakuba* females from allopatric populations. To understand why reinforced gametic isolation is confined to areas of secondary contact and has not spread throughout the entire *D. yakuba* geographic range, we studied the costs of reinforcement in *D. yakuba* using a combination of natural collections and experimental evolution. We found that *D. yakuba* males from sympatric populations sire fewer progeny than allopatric males when mated to allopatric *D. yakuba* females. Our results suggest that the correlated evolution of male and female reproductive traits in sympatric *D. yakuba* have associated costs (i.e., reduced male fertility) that prevent the alleles responsible for enhanced isolation from spreading outside the hybrid zone.

## INTRODUCTION

Reinforcement is the evolutionary process through which prezygotic reproductive isolation is strengthened by natural selection acting against the production of maladapted, infertile, or inviable hybrids (Dobzhansky 1937, Coyne and Orr 2004, Pfennig and Pfennig 2012). Reinforcement is expected to drive the evolution of prezygotic isolation in regions where two closely related species overlap and hybridize (Dobzhansky 1937, Coyne and Orr 2004). This generates a pattern of “reproductive character displacement,” in which individuals of different species found in the same area (sympatry) display greater behavioral isolation from one another than individuals from different areas (allopatry; Brown and Wilson 1956, Rice and Pfenning 2008, Pfennig and Pfennig 2009). Reinforcement is abundant and can be found in a wide range of taxonomic groups, including fungi (Anderson et al. 1980, Turner et al. 2010, Murphy and Zeyl 2015), animals (Noor 1995, Gerhardt 1994, Pfennig 1998, Rundle and Schluter 1998, Higgie et al. 2000, Haavie et al. 2004, Jaenike et al. 2006), and plants (Kay and Schemske 2008, Hopkins and Rauscher 2011, 2012, Hopkins 2013). Therefore, reinforcement could be common during the ‘completion’ of speciation (Hudson and Price 2014).

Data describing specific aspects of the process of reinforcement are becoming widespread: we now have precise measurements of the strength of reinforcing selection (Hopkins et al. 2014), know that reinforcement can occur despite gene flow (Sanderson 1989, Servedio 2000, Nosil et al. 2003 Matute 2010b), and have evidence that reinforcement can promote divergence within species, fostering speciation (Liou and Price 1994, Nosil 2005, Pfennig and Rice 2014, Nosil and Hohenloe 2012). Still, important aspects of reinforcement remain unexplored. Particularly we lack information regarding how variation in reproductive isolation is maintained across a species’ geographic range. The persistence of different levels of reproductive isolation between populations of a single species in the face of gene flow and stabilizing selection (Sanderson 1989, Servedio and Kirkpatrick 1997, Servedio and Noor 2003) poses a conundrum. One possible explanation is that reinforced reproductive isolation (RRI) favors phenotypes that are selected against in allopatry. In this case, the traits and alleles involved in reinforcement would not spread across a species’ geographic range but would remain confined to areas of secondary contact (Walker 1974, Caisse and Antonovics 1978, Howard 1993, Pfennig and Pfennig 2009).

Recent work suggests that phenotypes favored by reinforcing selection in sympatry are often disadvantageous in allopatry (Higgie and Blows 2008; Lemmon 2009; Hopkins et al. 2014; Pfennig and Rice 2014; Kozak et al. 2015). These studies provide evidence that reinforcing selection acting in sympatry can affect levels of reproductive isolation observed among conspecific populations: initiating the evolution of reproductive isolation and speciation through a process referred to as ‘cascading reinforcement’ (Howard 1993; Lemmon 2009; Kozak et al. 2015). To date, all reported cases of cascading reinforcement or of fitness costs associated with phenotypes involved in RRI come from studies in which premating reproductive isolating mechanisms are reinforced. For premating reproductive traits, it may be difficult to determine whether reproductive character displacement in sympatry results from indirect selection against the production of hybrids or direct selection on traits involved in species recognition (Shaw and Mendelson 2013).

In addition to traits involved in premating isolation, reinforcing selection can act on post-mating traits (Coyne 1974, e.g., gametic isolation: Lee et al. 1995, Wullschleger et al. 2002, Springer and Crespi 2007, Turner et al. 2010, Matute 2010 reviewed in Palumbi 2008). Examples of post-mating traits include proteins involved in interactions between reproductive tracts and gametes (reviewed in Palumbi 2008). Postmating-prezygotic phenomena have physiological and behavioral consequences that can influence both female and male fitness (Pitnick et al 1991, Yapici et al. 1998, Knowles et al. 2004, Markow and O’Grady 2008). Coevolution of these traits can be rapid when there is sexual conflict: ejaculate traits that directly increase male fitness can have deleterious effects on female fitness (Bateman 1948, Fowler and Partridge 1989, Chapman et al. 1995, Arnqvist and Rowe 2002, Wigby et al. 2005, Wigby et al. 2009). Previous experiments in *Drosophila* have demonstrated ongoing coevolution between the female reproductive tract and male ejaculate (Rice 1996, Holland and Rice 1999, Knowles and Markow 2001, Bono et al. 2011, Kelleher et al. 2011, Manier et al. 2013). Traits associated with the female reproductive tract and male ejaculate can therefore become finely tuned within populations (through continual antagonistic coevolution) but not necessarily across subpopulations within a species (Rice and Hostert 1993, Rice 1996, Manier et al. 2013). This provides a hypothetical mechanism for how RRI that evolves in sympatry could have cascading effects on levels of reproductive isolation among conspecific populations.

In this study we use the drosphilid flies *Drosophila yakuba* and *D. santomea* to explore the evolutionary dynamics of reinforcement in *D. yakuba. Drosophila yakuba* is a human-commensal species that is widespread throughout sub-Saharan Africa and has extended its range to islands in the Gulf of Guinea. On the volcanic island of São Tomé (off the coast of Cameroon and Gabon), *D. yakuba* occurs at low elevations (below 1,450 m), and is mostly found in open and semi-dry habitats commonly associated with human settlements (Lemeunier et al. 1986, Matute et al. 2009). In contrast, *D. santomea*, the sister species of *D. yakuba*, is endemic to the highlands of São Tomé where it is thought to exclusively breed on endemic figs (*Ficus chlamydocarpa fernandesiana*; Lachaise et al. 2000, Llopart et al. 2005a, b). The two species come into secondary contact and hybridize in the midlands of the mountain Pico de São Tomé within a well-defined hybrid zone (Llopart et al. 2005a, b). *Drosophila yakuba* females from this hybrid zone show higher postmating-prezygotic isolation toward males of *D. santomea* than do *D. yakuba* females from outside the hybrid zone (Matute 2010a), via an unknown mechanism acting within the female reproductive tract (Matute 2010a). There is no indication of RRI in *D. santomea*, even though some genetic variance seems to segregate in this species (Matute 2010b).

The RRI observed in sympatric *D. yakuba* can also evolve rapidly in experimental populations derived from allopatric lines (Matute 2010a, b). Matute (2010a, b) imposed strong selection against hybrids in experimental populations where *D. yakuba* and *D. santomea* were maintained in sympatry and both behavioral and gametic isolation increase in less than ten generations. These results show that the genetic variation required for behavioral and gametic isolation to evolve is present in allopatric populations. Experimental evolution can thus allow us to test factors that affect the reinforcement of reproductive isolation.

Here, we aim to determine whether *D. yakuba* lines that show RRI in female gametic isolation evolve correlated phenotypes in males, as would be predicted under a scenario of coevolution between female and male traits. In particular, we studied whether male fertility varies across an altitudinal transect spanning the range of *D. yakuba* on Pico de São Tomé. We show that natural lines of *D. yakuba* where females have RRI towards *D. santomea* males also have lower average male fertility in conspecific matings. Additionally, we show that when enhanced gametic isolation evolves after experimental sympatry with *D. santomea*, lower male fertility also evolves as a correlated trait. For *D. yakuba* females, enhanced postmating-prezygotic isolation from *D. santomea* males is advantageous in the hybrid zone, but our results suggest that lower male fertility outside of areas of potential interbreeding with *D. santomea* is a correlated cost of reinforced gametic isolation.

## METHODS

### Characterization of the hybrid zone

We collected males from the *yakuba* species subcomplex (*D. yakuba, D. santomea* and their reciprocal hybrids) in a transect on the north side of the island of São Tomé starting at sea level and ending at the *D. yakuba/D. santomea* hybrid zone at 1,200m above the sea level. The collection sites are shown in Supplementary Figure 1. We sampled *Drosophila* males from 17 localities as described in Matute (2015). We counted the number of *D. santomea*, *D. yakuba*, and both types of reciprocal F_1_ hybrids. To quantify levels of hybridization at each locality, we qualitatively scored the abdominal pigmentation of all the collected individuals that belonged to the *yakuba* subcomplex of species. We focused only on male flies, as they are much easier to identify than females. *Drosophila yakuba* males have a dark abdomen, while *D. santomea* males have a yellow abdomen. F_1_ hybrid males from the ♀*D. yakuba* ×♂ *D. santomea* cross have an intermediate abdominal pigmentation that is different from pure *D. yakuba* in the A3 and A4 segments (Llopart et al. 2002). F_1_ hybrid males from the reciprocal ♀*D. santomea* ×♂ *D. yakuba* cross have a patch of brown pigmentation in the A2 and A3 segments (Llopart et al. 2002). Counting F_1_ males constitutes a conservative estimate of the production of hybrids as it does not aim to identify individuals from advanced intercrosses.

**FIGURE 1.**
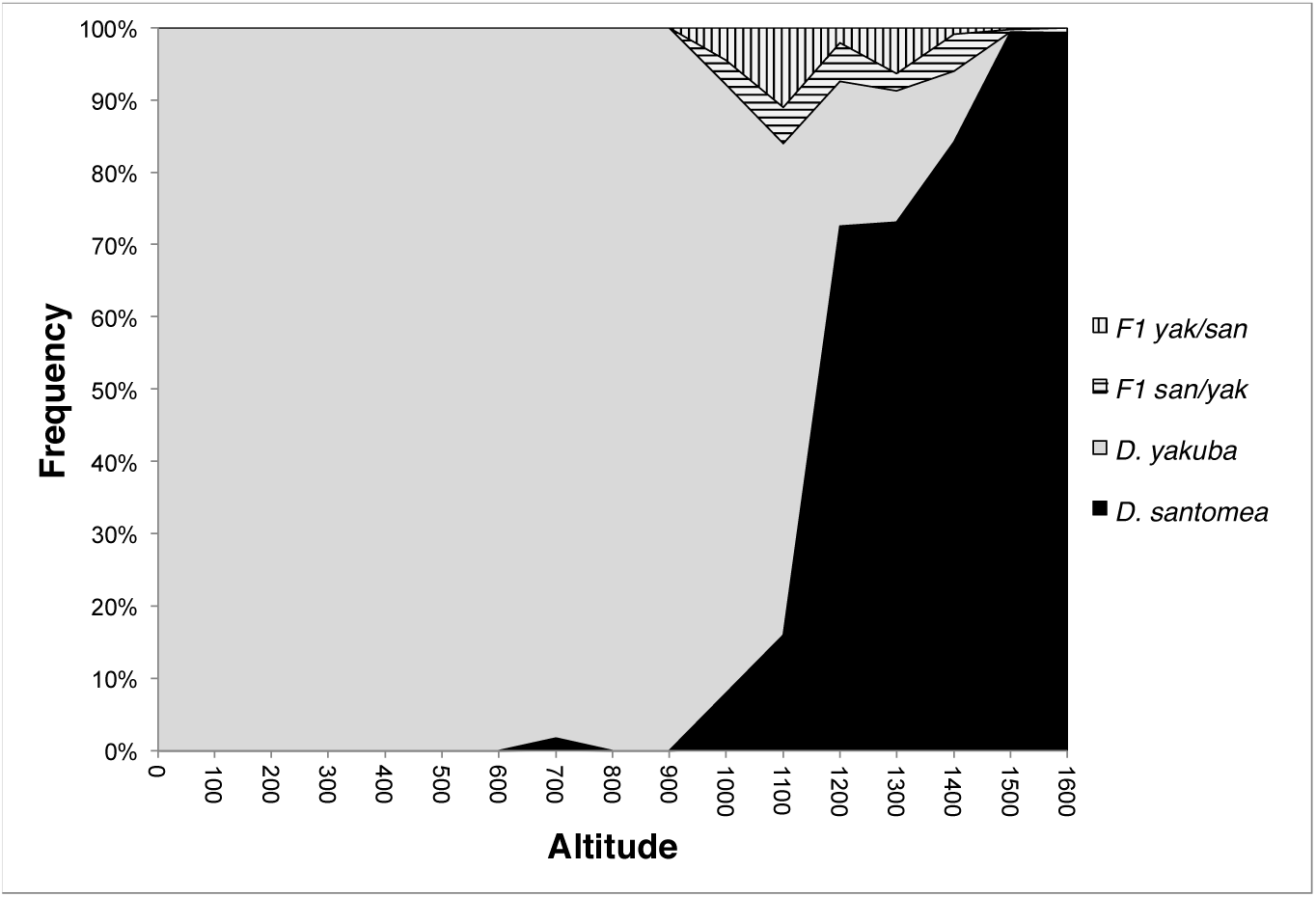
Abundance of *D. yakuba*, *D. santomea* and their F_1_ hybrids along the altitudinal gradient on Pico de São Tomé. Results presented are for F_1_ males that were identified based on their abdominal pigmentation.

### Stocks

In addition to collecting hybrid males, we established 344 isofemale lines from putative *D. yakuba* females collected from 17 different locations along the transect. Although *Drosophila yakuba*, *D. santomea,* and their hybrids have different colored abdomens, the differences between hybrid females and pure species are subtle (Llopart et al. 2002) and abdomen pigmentation varies within *D. yakuba* (Matute and Harris 2013). Therefore, we did not use abdominal pigmentation to distinguish pure species from hybrid females. Instead, we took advantage of the fact that pure species *D. yakuba* females produce only sterile males when crossed to *D. santomea* males (Lachaise et al. 2000) to determine if the collected females, and thus the isofemale lines, were pure species *D. yakuba* or advanced generation *D. yakuba* backcrosses. If the crossed females had ‘pure’ *D. yakuba* genotypes, then they would produce 100% fertile progeny when crossed with *D. yakuba* and 100% sterile progeny when crossed to *D. santomea.* On the other hand if putatively pure *D. yakuba* were in fact of recent hybrid origin, they would produce sterile male progeny when crossed to either parental (e.g. ∼90% if the female is a F_1_ hybrid; see SI for details). Of the 344 collected isofemale lines, we were able to successfully establish 297 isofemale stocks that fulfilled the criteria of being considered ‘pure’ *D. yakuba* (SI; Supplementary Table 1).

For experimental evolution assays (see below), we generated genetically heterogeneous strains of each species (i.e., synthetic lines) by combining virgin males and females from several isofemale lines collected on São Tomé outside of the hybrid zone (i.e., all lines were from allopatric populations). The synthetic *D. santomea* line (*D. santomea* SYN 2005) was generated by J.A. Coyne by combining six isofemale lines collected in 2005 at the field station Bom Sucesso (elevation 1,150 m). The synthetic *D. yakuba* line (*D. yakuba* SYN2005) was generated by combining five isofemale lines collected by J.A. Coyne in 2005 on Pico de São Tomé (elevation 880m). All stocks were kept in large numbers after they were created. All rearing was done on standard cornmeal/Karo/agar medium at 24°C under a 12 h light/dark cycle.

### Male fertility: sympatry versus allopatry

We quantified levels of male fertility by counting the number of progeny produced following crosses between females and males of 20 isofemale lines (all pairwise combinations). To ensure that these lines were unambiguously sympatric or allopatric ten were collected at the low elevation end of the transect described above (low elevation = allopatric lines) while the other 10 were collected in the hybrid zone (high elevation = sympatric lines). These 20 lines showed no evidence for adaptation to different temperatures (temperature covaries with elevation) or the evolution of reproductive isolating mechanisms apart from those described below (see SI for details). Moreover, these lines show only moderate genetic differentiation (median F_ST_ = 0.0503) and no evidence of large chromosomal differences (see SI for details). These results indicate that allopatric and sympatric lines differ primarily in whether they occurred in in the absence or presence of *D. santomea*.

For each of the twenty lines, we collected virgin males and females under CO_2_ anesthesia and kept them in isolation for 3 days in single-sex groups of 20 flies. On day 4, we conducted no-choice mating trials as previously described (Coyne et al. 2002, Matute and Coyne 2010). Briefly, we combined a single female and a single male, observed whether the pair mated and, if so, recorded copulation latency and copulation duration. Females that showed an abnormally short copulation (< 20 minutes) were discarded as no sperm transfer occurs before that time (Chang 2004). After 1 hour, we ended the observations and discarded any females that had not mated. To prevent females from remating, males were removed from each vial by aspiration after mating was finished. Each mated female was allowed to oviposit for 24 h. We then transferred the female to a fresh vial and counted the total number of eggs laid. The counting was repeated daily for 10 days. We scored 10 males per cross for a total of 4,000 males. Crosses were classified as being one of six possible types: ♀allopatric × ♂sympatric, ♀sympatric × ♂allopatric, ♀allopatric × ♂ allopatric (different isofemale lines), ♀ sympatric × ♂ sympatric (different isofemale lines), ♀allopatric × ♂ allopatric (within isofemale lines), and ♀ sympatric × ♂ sympatric (within isofemale lines).

We analyzed the data using a generalized linear mixed effect model (GLMM) in which the number of eggs laid by a female (fertility) was the response, the cross type was the fixed effect (the six cross types defined above), and the identity of the isofemale lines were random effects (female_line_ and male_line_). We used the R package ‘lme4’ (function ‘glmer’, Bates et al. 2011) and fitted the model, assuming Poisson distributed error, with the form:

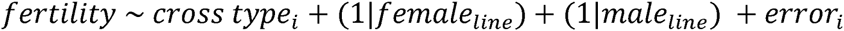

To determine whether the type of cross explained a significant proportion of variation in fertility, we compared the proportion of residual deviance explained by the model described above to one lacking the fixed effect of cross type using two methods: 1) a likelihood ratio test (‘lrtest’ function of the ‘lmtest’ R package) and 2) parametric bootstrapping (‘PBmodcomp’ function in the ‘pbkrtest’ R package, Halekoh and Højsgaard 2014). Parametric bootstrapping was carried out using 100 bootstrapped samples. We also used pairwise Tukey contrasts (‘glht’ and ‘linfct’ functions of the ‘multcomp’ R package) to determine the types of crosses, if any, differed in level of fertility.

### Geographic distribution of gametic isolation and male fertility

#### i) Female gametic isolation from *D. santomea*

When mated to *D. santomea* males, sympatric *D. yakuba* females (i.e., from the hybrid zone) lay fewer eggs than allopatric females (Matute 2010a). We quantified the geographic scale of female gametic isolation in *D. yakuba* isofemale lines collected along the altitudinal transect across São Tomé. We used ten lines from each of ten collection locations for a total of 100 isofemale lines. Lines were chosen randomly from the 297 pure *D. yakuba* lines and are listed in Supplementary Table 1. We collected virgin *D. yakuba* females from each of these 100 isofemale lines and let them age to four days (as described above) and then mated them to *D. santomea* SYN2005 males (heterospecific cross). Mated females were kept, and the number of eggs they produced was scored every 24 hours over the course of ten days. In parallel, we mated *D. yakuba* females from each isofemale line to males from the same isofemale line and counted the number of eggs produced by each female (conspecific cross). Heterospecific and conspecific pairings were monitored in parallel to ensure that mating occurred under the same environmental conditions. For each line, we scored the number of eggs produced by fifteen females in both heterospecific and conspecific matings (N =15 females × 100 lines = 1,500 females for each type of mating). The proportion of eggs produced after heterospecific matings relative to conspecific matings was taken as an inverse proxy for the magnitude of gametic isolation (i.e., the more eggs produced after a single heterospecific mating the weaker the gametic isolation; Chang 2004).

To analyze whether there were differences in the strength of gametic isolation between isofemale lines collected from sympatric versus allopatric regions, we fitted GLMMs with Poisson distributed error using the “glmer” function in the “lme4” R package. We treated the number of eggs produced in heterospecific and conspecific crosses as separate data sets. The two GLMMs were therefore constructed as:

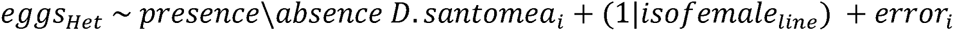

and

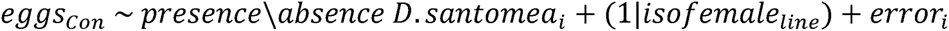

We assessed significance of the presence \ absence of *D. santomea* on the number of eggs produced using both maximum likelihood tests (LRT, 1 degree of freedom) and parametric bootstrapping (100 bootstrap samples, function ‘PBmodcomp’ in the ‘pbkrtest’ R package; Halekoh and Højsgaard 2014) that compared the two models to two ‘null’ models that lacked the fixed effect of presence \ absence of *D. santomea.*

#### ii) Intraspecific male fertility

We next quantified male fertility along the altitudinal transect using the same 100 *D. yakuba* isofemale lines used to study female gametic isolation. We crossed males from each of these lines with conspecific females from two tester stocks: one allopatric and one sympatric. *Drosophila yakuba* Täi18 (hereafter referred to as allopatric_Täi18_), is an allopatric isofemale line, collected in the Täi forest on the border between Liberia and Ivory Coast. BOSU1250.5 (hereafter referred to as sympatric_BOSU1250.5_) is a *D. yakuba* line collected in 2005 at the heart of the São Tomé hybrid zone and is considered sympatric. We collected virgin males and when they were 4 days old, allowed them to mate to virgin females from either tester stock following the mating procedure described above (no-choice trials). The number of eggs produced over ten days was assessed as a proxy for male fertility with females from different populations. Crosses with allopatric_Täi18_ and sympatric_BOSU1250.5_ were considered different datasets. Each dataset was analyzed by fitting a GLMM with Poison distributed error where the number of eggs produced per cross was the response, the origin of the male (whether the isofemale line is sympatric or allopatric) was the fixed effect, and isofemale line was a random effect:

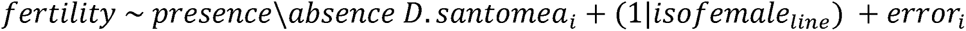

For both ‘allopatric’ and ‘sympatric’ data sets, the model described above was compared with a model without the fixed effect of presence \ absence of *D. santomea* using both maximum likelihood tests (LRT, 1 degree of freedom) and parametric bootstrapping (100 bootstrap samples, function ‘PBmodcomp’ in the ‘pbkrtest’ R package; Halekoh and Højsgaard 2014).

### Correlation between gametic isolation and male fertility

To determine whether the average magnitude of gametic isolation from *D. santomea* males and average male fertility per isofemale line were correlated across isofemale lines, we used Spearman’s rank correlation tests implemented in the ‘stats’ package in R (function ‘cor.test’). Bonferroni correlation tests gave similar results.

### Experimental Sympatry

Previous work has shown that strong selection against hybridization can drive the evolution of enhanced gametic isolation in female *D. yakuba* that are kept in experimental sympatry with *D. santomea* (Matute 2010a,b). If reinforced gametic isolation and reduced male fertility are correlated traits, they should evolve together in experimental sympatry. We took an experimental evolution approach to test this hypothesis and establish whether lower male fertility between allopatric and sympatric populations of *D. yakuba* could evolve as a correlated trait with enhanced gametic isolation from *D. santomea* observed in females.

We kept 23 populations (bottles) of *D. yakuba*, originally derived from a synthetic allopatric population (*D. yakuba* SYN2005) in experimental sympatry with *D. santomea* (*D. santomea* SYN2005) for ten generations following previously described experimental protocols (Koopman 1950, Higgie et al. 2000, Blows and Higgie 2002, Matute 2010a). Each bottle contained 250 *D. yakuba* females, 250 *D. yakuba* males, 250 *D. santomea* females, and 250 *D. santomea* males. To set up each successive generation, we collected 250 flies of each sex of *D. yakuba* (identifiable by their abdominal pigmentation) as virgins from the experimental bottles and transferred them into new bottles. To reconstitute sympatric conditions, 250 *D. santomea* flies of each sex (collected as virgins from stock bottles) were added to each bottle. All flies were collected between seven and ten hours after eclosion once the flies had already achieved their adult pigmentation. Hybrids were recognized by their abdominal pigmentation (intermediate – yet different – between the two parental species) and were discarded. This procedure was followed for ten generations. Twenty-three control populations (i.e. bottles) of *D. yakuba* were maintained in parallel with the same number of conspecifics (500 flies per bottle) but in the absence of *D. santomea*. The maintenance conditions and population size of *D. yakuba* were the same between the experimental sympatry and control bottles. The strength of gametic isolation was measured after ten generations of experimental sympatry using methods described previously (see “Geographic distribution of gametic isolation and male fertility” above).

We compared levels of female gametic isolation from *D. santomea*, male fertility with sympatric_BOSU1250.5_ females, and male fertility with allopatric_Täi18_ females after 10 generations of experimental sympatry. We fitted generalized linear models (GLMs) with Poisson distributed error (i.e., Poisson regression) in which the magnitude of gametic isolation or male fertility was the response and the generation (0, or 10) was the fixed effect. To assess the affect of time (i.e., generation) on the evolution of gametic isolation or male fertility we used likelihood ratio tests comparing models including versus excluding this term. Models were fitted using the ‘glm’ function in R. Finally, we used Spearman’s rank correlation to compare whether there was a correlated response in levels of isolation from *D. santomea* observed in females and isolation from allopatric *D. yakuba* observed in males across experimental lines.

## RESULTS

### Characterization of the hybrid zone

We sampled from across the hybrid zone between *D. santomea* and *D. yakuba* on the island of Pico de São Tomé to measure the distributions of *D. santomea*, *D. yakuba,* and their two reciprocal hybrids. The abundances of each of the four genotypes (2 pure species and two F_1_ hybrids) at 17 locations along the altitudinal transect is shown in Figure 1. Hybrids were found in the same locations as four years prior (Llopart et al. 2005a), suggesting that the hybrid zone has remained stable since it was first reported. Notably we found a few *D. santomea* males at 700 m indicating that *D. santomea* males sometimes wander out of the canonical distribution previously reported for the species.

### Male fertility: Sympatry versus allopatry

We have previously described the reproductive advantage obtained by *D. yakuba* females that evolved RRI both in nature, and under experimental evolution (Matute 2010a, 2010b). We hypothesized that there may be an associated cost to RRI that prevents it from spreading to allopatric populations of *D. yakuba*. We mated males and females of allopatric and sympatric lines in all 4 possible combinations, and counted how many eggs were produced by each mating type. We found that males from sympatric areas sire fewer progeny than allopatric males when mated to allopatric females but not when mated to sympatric females (Figure 2b). Generalized linear mixed models fitted to the data showed that the type of cross being conducted (the fixed effect in these models) explained a significant proportion of residual deviance in the number of eggs produced when compared to a null model lacking this term (LRT: χ^2^ = 700.09, df = 5, *P* < 1 × 10^-15^; parametric bootstrapping: *P* = 0.01; Supplementary Table 3). Pairwise comparisons revealed that the average number of eggs laid in ♂ sympatric x ♀ allopatric crosses (mean = 75.38, standard error [SE] = 0.21) was significantly lower compared to the number of eggs laid in all other cross types (range of mean [and SE] across other types of cross = 88.79 – 90.23 [0.28 – 0.91]); all *P* < 0.0001; Figure 1; Supplementary Table 4).

**FIGURE 2.**
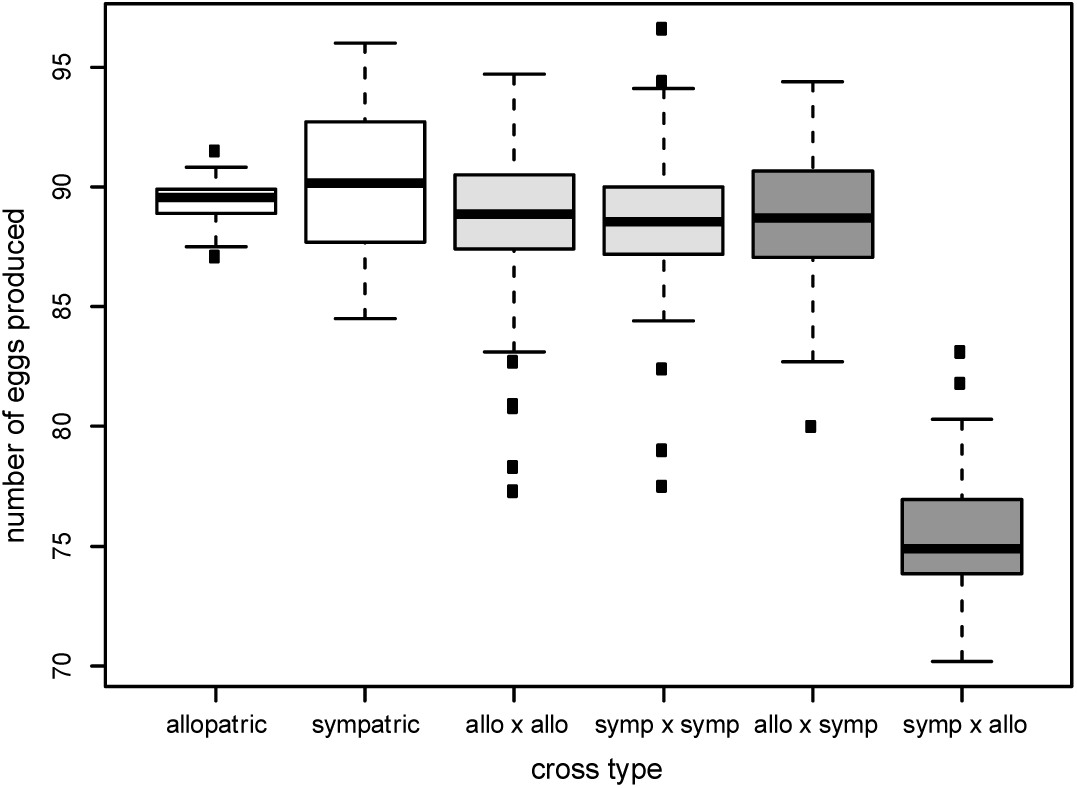
Males from a sympatric population show reduced fertility when mated to allopatric females. We crossed males and females from 20 isofemale lines in all possible combinations and counted the number of eggs produced over ten days Crosses were split into six categories depending on the identity of the female and the male involved in the cross. The y-axis shows the average number of eggs produced after 10 days across crosses in each category. Categories of crosses are (from left to right): males and females from the same allopatric isofemale line (white box), males and females from the same sympatric isofemale line (white box), ♂ sympatric x♀ sympatric from different isofemale lines (light grey), ♂allopatric × ♀ allopatric from different isofemale lines (light grey), ♂ allopatric × ♀ sympatric lines (dark grey), and ♂sympatric ×♀ allopatric lines (dark grey) lines. Sympatric males produced fewer progeny when mated to allopatric females than any of the other possible crosses.

### Geographic distribution of gametic isolation and male fertility

We characterized the geographic distribution of RRI along an altitudinal transect on Pico de São Tomé by collecting 10 *D. yakuba* isofemale lines at each of 10 different sites along the transect (N=100, including the 20 lines we used above) and measuring the strength of gametic isolation in *D. yakuba* females towards *D. santomea* males. We focused on lines that were putatively pure *D. yakuba* (rather than lines with potentially admixed ancestry; see SI). We fitted GLMMs using sympatry with *D. santomea* as a fixed effect and isofemale line as a random effect. We found that the magnitude of gametic isolation between populations of *D. yakuba* and *D. santomea* is affected by sympatry with *D. santomea* (LRT: χ^2^ = 54.37, df = 1, *P* = 1.66 × 10^-13^ ; parametric bootstrapping: *P* = 0.0099; Figure 3a, black boxes), conforming the expectation that enhanced gametic isolation is driven by maladaptive hybridization with *D. santomea*. Female fertility in conspecific matings also differed between allopatric and sympatric lines (LRT: χ^2^ = 86.97, df = 1, *P* < 1 × 10^-15^; parametric bootstrapping: *P* = 0.0099; Figure 3a, grey boxes); however, the magnitude of the difference in mean number of eggs produced between sympatric and allopatric *D. yakuba* females was much lower in conspecific crosses (mean eggs [SE]: sympatric females = 86.70 [0.17]; allopatric females = 93.19 [0.18]) when compared to heterospecific crosses (mean eggs [SE]: sympatric females = 34.60 [0.47]; allopatric females = 63.35 [0.60]).

**FIGURE 3.**
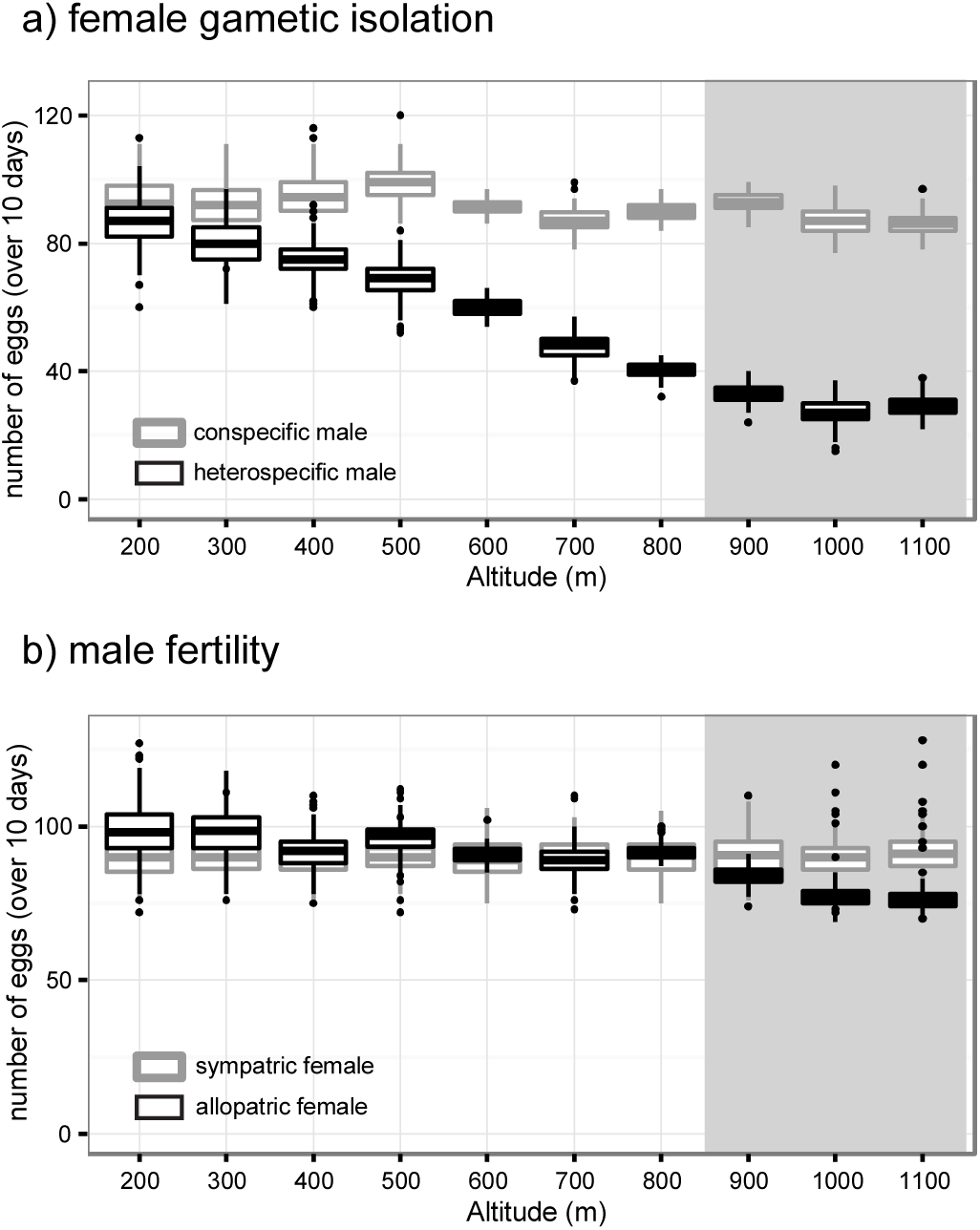
Levels of gametic isolation and male fertility across the altitudinal transect. We measured the levels of gametic isolation and male fertility in 100 lines collected along the altitudinal transect in São Tomé (10 lines per elevation / site). Gray shaded polygons to the right of each panel demarcate the location of the hybrid zone. a) There is no gametic isolation among females when crossed to *D. yakuba* males (conspecific crosses; gray boxes); however, isofemale lines collected from closer to the hybrid zone have lower fertility (i.e. higher gametic isolation) when crossed to *D. santomea* (heterospecific crosses, black boxes). b) Male fertility, measured as the number of eggs produced, is lower for isofemale lines collected from the hybrid zone when mated to female *D. yakuba* from an alloparic isofemale line (black boxes), but not when mated to female *D. yakuba* from a sympatric isofemale line (gray boxes). Males from sympatric populations showed lower fertility when mated to allopatric females, but not when mated to sympatric females.

We next asked whether male fertility varied among the same lines collected along the transect. We measured male fertility by mating males from each isofemale line to either allopatric_Täi18_ females or sympatric_BOSU1250.5_ females (N=15 crosses per tester line). These two lines had previously been identified as representative sympatric and allopatric lines (Matute 2010a). The results from these crosses are shown in Figure 3b. We fitted two GLMMs similar to those used to study the magnitude of gametic isolation observed in females: one for matings with sympatric_BOSU1250.5_ females and one for matings with allopatric_Täi18_ females. Male fertility did not change between regions of sympatry and allopatry when mated to sympatric_BOSU1250.5_ females (LRT: χ^2^ = 86.97, df = 1, *P* = 1 × 10^-15^; parametric bootstrapping: *P* = 0.0099; Figure 3b, grey boxes). By contrast, male fertility differed between allopatric and sympatric regions when males were mated to allopatric_Täi18_ females (LRT: χ^2^ = 0.2144, df = 1, *P* = 0.64; parametric bootstrapping: *P* = 0.63; Figure 3b, black boxes). The number of eggs produced by allopatric females suggests that male fertility is lowest in lines that are sympatric with *D. santomea* (Figure 3b) and that the geographic distribution of reduced male fertility on São Tomé mirrors that of enhanced gametic isolation.

### Correlation between gametic isolation and male fertility

We also examined whether the magnitude of female gametic isolation was correlated with male fertility in crosses made with allopatric_Täi18_ and sympatric_BOSU1250.5_ females. We evaluated the correlation between the average strength of gametic isolation and average male fertility when mated with allopatric and sympatric females using 25 females and 25 males per isofemale line. Female gametic isolation was correlated with male fertility with allopatric_Täi18_ females (Spearman’s rho = 0.885, *P* < 1×10^-10^; Figure 4a) but not with sympatric_BOSU1250.5_ females (Spearman’s rho = 0.116, *P* = 0.250; Figure 4b). These results illustrate that female gametic isolation toward heterospecifics in sympatry might evolve at the cost of reduced male fertility in conspecific allopatric matings.

**FIGURE 4.**
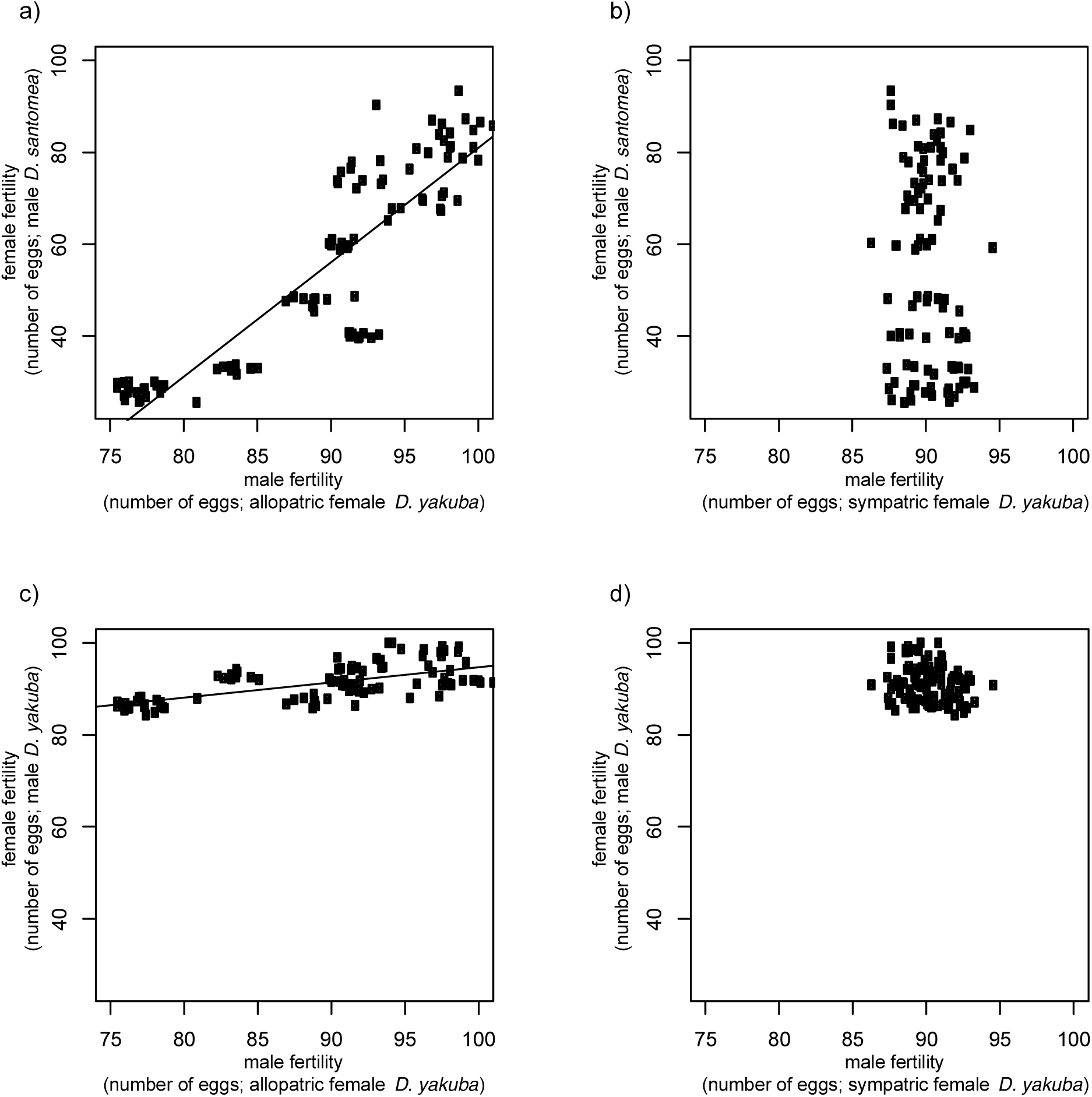
Correlations between male fertility and female gametic isolation in natural populations. a) Male fertility in matings with allopatric females is correlated with the number of eggs laid by females (female gametic isolation) after heterospecific matings with *D. santomea* (as an inverse proxy of the strength of gametic isolation) (Spearman’s rho = 0.885, P < 1 ×10^-10^). Male fertility is not correlated female fertility when mated with sympatric females (Spearman’s rho = 0.116, P = 0.250). c) The number of eggs laid by females after conspecific matings is also correlated with male fertility in matings with allopatric females (Spearman’s rho = 0.605, P **<**1 ×10^-10^), but not with male fertility in matings with sympatric females (d; Spearman’s rho = -0.149, P = 0.140). Lines shown only for significant correlations.

### Experimental Sympatry

Previous experiments have shown that female *D. yakuba* can evolve increased gametic isolation from *D. santomea* males when evolved under sympatric conditions (Matute 2010a, b). We repeated this experiment and again found that *D. yakuba* females laid fewer eggs when mated to heterospecific *D. santomea* males following 10 generations of sympatry than were laid at generation 0 (Poisson GLM, LRT: χ^2^ = 2767.7; df = 1; *P* < 1×10^-10^, Figure 5a). The number of eggs laid following conspecific matings remained the same (LRT: χ^2^ = 2.43; df = 1; *P* = 0.12; Figure 5a). We next looked at whether reduced conspecific male fertility with both allopatric_Täi18_ and sympatric_BOSU1250.5_ *D. yakuba* evolved as a correlated trait with increased female gametic isolation from *D. santomea*. We found no difference in male fertility among the experimentally evolved lines when mated to sympatric_BOSU1250.5_ females (LRT: χ^2^ = 1.06; df = 1; *P* = 0.304; Figure 5b). However, male fertility in matings with allopatric_Täi18_ females showed a significant decrease over ten generations of experimental sympatry (LRT: χ^2^ = 743.9; df = 1; *P* < 1×10^-10^, Figure 5b). In control populations of *D. yakuba* raised in parallel to the experimental populations but with *D. santomea* absent, we observed no change in female gametic isolation (LRT: χ^2^ = 0.54; df = 1; *P* = 0.462), or male fertility (LRT: χ^2^ = 0.89; df = 1; *P* = 0.347) over the same 10 generations of evolution. Finally, we observed a significant correlation between levels of female gametic isolation from *D. santomea* and male fertility with allopatric_Täi18_ females across experimental replicates (Spearman’s rho = 0.69; *P* < 1 × 10^-10^; Figure 5c) but not with male fertility in matings with sympatric_BOSU1250.5_ females (Spearman’s rho = -0.046, *P* = 0.115). These results provide experimental evidence that when gametic isolation from *D. santomea* males evolves via reinforcing selection acting on *D. yakuba* females, there is a correlated cost of postmating prezygotic incompatibility between sympatric males and conspecific females from allopatric populations.

**FIGURE 5.**
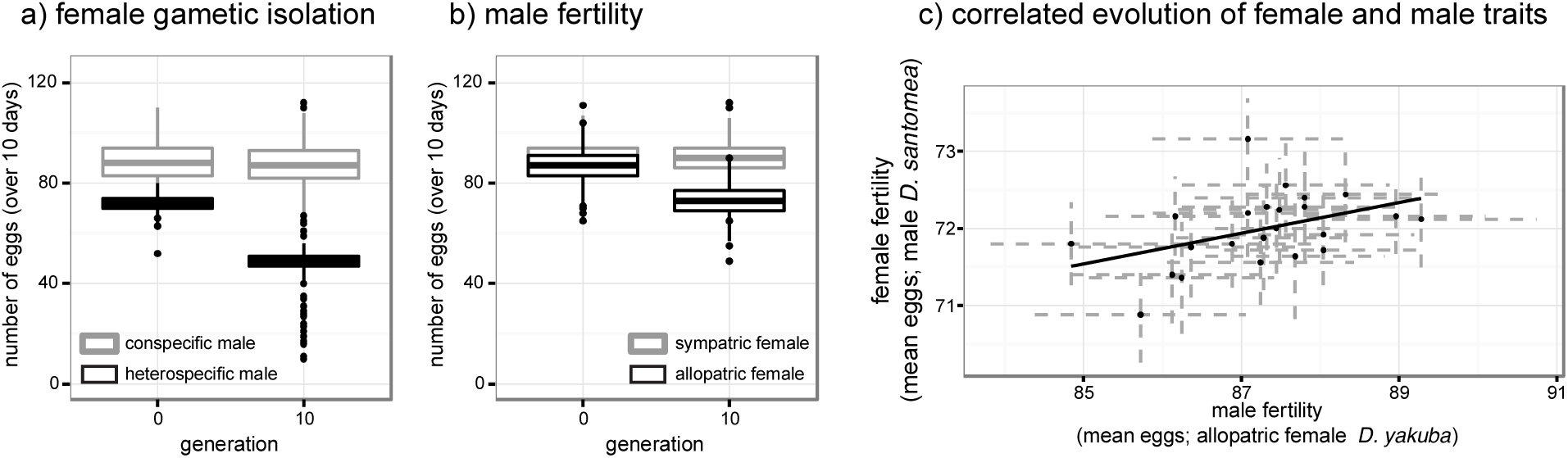
Experimental evidence that gametic isolation and decreased male fertility coevolve after experimental sympatry with *D. santomea*. a) After 10 generations of experimental sympatry, allopatric *D. yakuba* lines evolved enhanced gametic isolation (black boxes) with no change in conspecific fertility (gray boxes). b) The same evolved lines also showed a decrease in their male fertility when males are mated to an allopatric line of *D. yakuba* (black boxes) but no decrease in their fertility when mated to females from a sympatric line of *D. yakuba* (gray boxes). c) Within experimental populations, the degree of female gametic isolation that evolved over 10 generations (“female fertility”; smaller values represent higher levels of isolation) was correlated with levels of male fertility with allopatric *D. yakuba* females. Points in “c)” represent means for each of 23 experimental populations and dashed gray lines represent ± one standard error. Control populations showed no change in gametic isolation from *D. santomea,* female fertility with conspecific males, male fertility when males are mated to allopatric *D. yakuba,* or male fertility when mated to sympatric *D. yakuba* (see text for statistics).

## DISCUSSION

We have used observation and experimental evidence to show that the reinforcement of gametic isolation between *D. yakuba* females and *D. santomea* leads to the correlated evolution of intraspecific postmating prezygotic isolation as the results of reduced conspecific male fertility. Our results suggest that there is ‘fine-tuning’ between male and female reproductive traits within populations of *D. yakuba.* Furthermore, we describe a cost associated with RRI in regions of sympatry with *D. santomea*: when sympatric *D. yakuba* males mate with allopatric *D. yakuba* females, they suffer from decreased fertility. Our finding provides a potential causal mechanism for the localized effect of reinforcing selection on reproductive isolation: alleles that underlie female traits that are adaptive in sympatry are associated with alleles that underlie male traits that are maladaptive in geographic regions outside of hybrid zones.

Although enhanced gametic isolation in females is highly beneficial in the presence of *D*. *santomea* (by reducing maladaptive hybridization) it is associated with conditionally lower male fertility. This correlated cost, coupled with the fact that enhanced gametic isolation provides no fitness benefit outside the hybrid zone, likely hampers the spread of the alleles underlying enhanced RRI into allopatry. Our results shed light on why the observed reinforced gametic isolation in the *D. yakuba/D. santomea* hybrid zone is confined to areas of secondary contact. Together, these results provide evidence for cascading reinforcement among populations of *D. yakuba* on the island of São Tomé.

### Alternate explanations for observed patterns of reproductive isolation

Reinforced reproductive isolation is one means by which natural selection can lead to the completion of speciation (Dobzhansky 1937, Lukhtanov et al 1995, Coyne and Orr 2004). However, it has remained unclear how the signature of reinforcement in hybrid zones persists despite migration between allopatric and sympatric populations (Walker 1974, Liou and Price 1994, Servedio and Noor 2003). Strong phenotypic differentiation between sympatric and allopatric populations can be generated through a variety of processes that do not involve selection against ‘reinforcing’ alleles in allopatric regions. For example, local adaptation to different environments, geographical structuring of populations, drift, and/or genetic incompatibilities could all contribute to reductions in gene flow among conspecific populations and lead to the incidental evolution of reproductive isolation. For the populations and lines of *D. yakuba* we focused on in this study, we addressed these issues and found it unlikely that phenotypic differentiation between allopatric and sympatric populations results from *i*) local adaptation to temperature, *ii)* forms of reproductive isolation other than male fertility, *iii*) strong genetic differentiation among populations at alternate ends of the altitudinal transect, or *iv*) chromosomal inversions limiting genomic admixture between demes (Hoffman et al. 2004, Kirkpatrick and Barton 2006) (see SI for details). These results are not surprising given that the collection sites for the hybrid zone and the allopatric areas are separated by less than 10 km; a distance some *Drosophila* species can travel overnight (Jones et al. 1981, Coyne et al. 1982, Coyne and Milstead 1987 but see Timofeef-Ressovsky and Timofeef-Ressovsky 1940; reviewed in Powell 1997). The fact that we observe some non-zero genetic differentiation among isofemale lines derived from females sample from opposite ends of the altitudinal transect suggests that reinforcing selection acting in sympatry – and selection acting in allopatry – could help drive genetic differentiation among conspecific populations: a supposition that warrants further investigation.

### Fitness tradeoffs and the cost of reinforcement

The reduced fertility we observed when sympatric males mated to allopatric females represents a fitness tradeoff between selection acting in regions where *D. yakuba* occurs in sympatry with *D. santomea* and selection acting in allopatry. Reduced male fertility also implies indirect costs to females with enhanced gametic isolation in sympatry because their sons will be less fit than those of allopatric females. The tradeoffs could explain why reinforced gametic isolation has not spread throughout the whole geographic range of *D. yakuba* on São Tomé. Our results add to the growing list of examples of traits favored through reinforcement in regions of sympatry but selected against in allopatry (Lemmon 2009; Hopkins and Rausher 2014; Pfennig and Rice 2014; Kozak et al. 2015). We do not mean to imply that reduced male fertility is the only cost associated with enhanced gametic isolation in *D. yakuba* females from the hybrid zone. It is possible that unexplored traits also hamper the spread of enhanced gametic isolation between sympatric and allopatric regions.

### Coevolution of female and male reproductive traits

At the phenotypic level, our findings highlight the tight link between the female reproductive tract and male ejaculate. In the case of *D. yakuba* we present here, male performance depends on the genotype of his mate (Figures 2, 3, 4 and 5). The evolution of gametic isolation towards *D. santomea* in female *D. yakuba* leads to a co-evolutionary change in male traits (manifested as reduced fertility with allopatric conspecifics). This result suggests that there is selectable genetic variation for female and male reproductive traits segregating within natural populations; a result seen in previous studies (Pitnick and Miller 2000, Miller and Pitnick 2002). A model of coevolution between male and female reproductive traits in *D. yakuba* would suggest that reinforcing selection can trigger changes in female traits that reduce the production of maladaptive heterospecific offspring, which in turn leads to coevolutionary changes in conspecific male traits.

Several lines of evidence indicate that the traits involved in the interactions between the ejaculate and the female reproductive tract are constantly diverging both between and within species (Manier et al. 2013). First, comparative studies have revealed correlated evolution of sperm and female reproductive tract morphology in *Drosophila* (Pitnick et al. 1997, 1999). Specifically, across the whole genus *Drosophila*, sperm length (among other sperm traits) is highly variable (Joly and Bressac 1994), and there is a strong positive relationship across species between the length of sperm and the length of the seminal receptacle, the main sperm-storage organ in the female (Pitnick et al. 1999). Furthermore, experimental evolution of *D. melanogaster* in the laboratory has revealed that increased seminal receptacle length can drive the evolution of increased sperm length (Pitnick and Miller 2000, Miller and Pitnick 2002). Our combination of comparative and experimental results suggest that coevolution between the sexes can drive correlated evolution between sperm and female reproductive tract traits, with evolution in female traits resulting in selection on corresponding traits in males.

At the molecular scale, components of the seminal fluid (such as the accessory gland proteins, Acps), are known to evolve faster than the rest of the genome (Ram and Wolfner 2007). Approximately 36% of Acps shared among members of the *D. melanogaster* subgroup (*D. melanogaster, D. simulans, D. sechellia, D. erecta,* and *D. yakuba*) appear to have evolved under the influence of positive selection (Swanson et al. 2001a; Mueller et al. 2005; Haerty et al. 2007). Additional studies have shown that Acps evolve rapidly within species (Begun et al. 2000; Kern et al. 2004; Begun and Lindfors 2005; Schully and Hellberg 2006). Such accelerated rates of molecular evolution are not unique to male proteins but have also been observed in genes related to fertilization and female receptors (Andres and Anrqvist 2001, Swanson et al. 2004, Lawniczak and Begun 2007, Chow et al. 2010). These results suggest that Acps and female receptors, the molecular underpinnings of male × female post-mating interactions, are prone to rapid evolution and species harbor sufficient genetic variation in these traits for natural and sexual selection to act upon.

### Caveats and conclusions

Our results come with at least three caveats. First, it is possible that sympatric females are of hybrid ancestry. We assessed this issue by measuring sterility of offspring when females were crossed to either *D. yakuba* and *D. santomea* (see SI for details). However, this assay will only reveal ‘hybrid’ *D. yakuba* lines that contain alleles that are involved in hybrid sterility with *D. santomea*; it is possible that other parts of the genome have admixed ancestry, although it is unclear how this might affect male fertility.

A second caveat comes from the population cage sizes used for experimental evolution. It has been shown that high population densities reduce fitness and influence behavior in a variety of organisms (e.g., Booth 1995, Zachar and Neiman 2013, Matute 2014). We chose to have cages with the same number of *D. yakuba* individuals, to control for levels of genetic variance between treatments; however, we did not explore the evolution of reproductive isolation at different experimental densities.

Thirdly, it is worth noting that even though reduced male fertility and enhanced gametic isolation are correlated traits, we have no evidence that these traits have the same genetic basis or that the alleles controlling the two traits are genetically linked. Since female receptivity × male ejaculate interactions are finely tuned, changes in female phenotypes, such as sperm retention, might cause concomitant changes in male traits related to ejaculate quality without the need to invoke genetic linkage. Future work using crosses will need to be done to further elucidate this connection.

It has long been argued that the evolution of reproductive isolation could be constrained by sexual selection, and that pleiotropic effects of reproductive isolation would either accelerate or hinder speciation (reviewed in Panhuis et al. 2001, Ritchie 2007). Our data demonstrate that the evolution of reinforcement is not free of associated costs, and that sexual selection might oppose reinforcing selection during speciation (Ritchie 2007, Safran et al. 2013). Other cases have been identified in which reinforcing selection leads to the evolution of traits unfavored by sexual selection (*Spea multiplicata*: Pfennig 1998; *Drosophila serrata*: Higgie et al. 2000).

Finally, our results contribute to the body of work demonstrating that locally adaptive traits can have unexpected costs that might hamper their expansion across the entire range of a species (Littlejohn and Loftus-Hills 1968; Zouros and D’Entremont 1980; Higgie and Blows 2008; Hopkins et al. 2014; Pfennig and Rice 2014; Kozak et al. 2015). When a trait contributes to reproductive isolation, the pattern of enhanced reproductive isolation in sympatry that is the hallmark of reinforcement can be explained by selection against the phenotypes involved outside of the hybrid zone.

